# Mapping DHPS Evolvability: Identification of Novel Evolutionarily Critical Sub-Structure through Evolutionary and Structural Analyses of DHPS

**DOI:** 10.1101/2024.10.16.618629

**Authors:** Dwipanjan Sanyal, A. Shivram, Suharto Banerjee, Vladimir N. Uversky, Aneesh Chivukula, Krishnananda Chattopadhyay, Sourav Chowdhury

## Abstract

Protein evolution shapes pathogen adaptation-landscape, particularly in developing drug resistance. The rapid evolution of target proteins under antibiotic pressure leads to escape mutations leading to the problem of antibiotic resistance. A deep understanding of the evolutionary dynamics of antibiotic target proteins presents a plausible intervention strategy for disrupting the evolutionary trajectory of resistance. Mutations in Dihydropteroate synthase (DHPS), an essential folate pathway protein and a key target for sulfonamide antibiotics, result in reduced antibiotic binding, leading to resistance. Deploying an array of statistical analyses on the DHPS sequence-space and integrating those with deep mutational analysis and structure-based network-topology models we identified critical DHPS-subsequences. Our analysis of the frustration landscape of DHPS predicts how conformational and mutational changes shift the energy distributions within the DHPS substructures. Combining dimensionality reduction and optimality analysis we identified a substructure critical to DHPS evolvability, and computed its druggablity. Our integrated evolution and structure-informed framework identified a DHPS-substructure with significant evolutionary and structural impact. Targeting this region could constrain DHPS evolvability and disrupt the resistome, presenting a new avenue for antibiotic development and contributing to the broader effort to address the problem of antibiotic resistance.

## Introduction

Protein evolution is a dynamic process driven by genetic variations that accumulate over time, altering amino acid sequences and, consequently, protein structure, function, and expression levels. This variability shapes adaptive responses, particularly in the context of pathogen virulence and the emergence of drug-resistant pathogens. The rapid adaptation of target proteins under antibiotic selection pressure poses a significant threat^1–3^, leading to escape mutations further conferring drug resistance^4–6^. Understanding the mechanisms underlying protein evolution and the acquisition of drug resistance is critical for developing effective strategies to combat this global health challenge. A more appropriate intervention would be disrupting the evolutionary trajectory of accumulating resistance, which requires a comprehensive understanding of the evolutionary dynamics influencing antibiotic target proteins^7^. Evolutionarily significant regions reflect functional constraints that limit protein evolvability and represent strategic therapeutic targets to mitigate antibiotic resistance by minimizing the possibility of the emergence of escape mutants^8^. Identifying these evolutionarily constrained regions is critical for developing evolution-proof interventions against the escalating threat of antibiotic resistance.

Our present study focuses on bacterial Dihydropteroate synthase (DHPS), an essential folate pathway protein^9^. Being an anti-folate target, DHPS is a prime example of how evolutionary processes drive drug resistance, given its critical role in folate biosynthesis and its frequent mutations in response to sulfonamide antibiotics. Folic acid is an essential precursor for the biosynthesis of purines, pyrimidines, and amino acids - building blocks for various cellular processes^10,11^. Multiple antibiotics target folate pathway proteins, such as folic acid, which is essential in cell division and helps in the production of nucleic acids^12,13^. Sulphonamides have been proven to be effective as broad-spectrum antimicrobials due to the ubiquitous presence of DHPS in lower organisms, coupled with its absence in higher organisms^14^. DHPS is a 267-amino acid enzyme encoded by the folP gene^15^, with a dimeric (β/α) TIM (Triose phosphate isomerase) barrel structure. It catalyzes the formation of 7,8-dihydropteroate from para-aminobenzoic acid (pABA) and dihydropterin pyrophosphate (DHPP) through an SN2 mechanism, a critical step in the folate pathway yielding tetrahydrofolate, essential for nucleotide synthesis^16^. The DHPS-active site, located at the C-terminal end of the barrel, includes a pterin binding site, an anion binding pocket facilitating PPi release, and flexible loops that bind pABA^15^. Active-site loop mobility is vital for forming the pABA binding pocket during the reaction. Sulfonamides, as competitive inhibitors, mimic pABA to bind the active site^17^, forming a metabolically inactive product.

The essential role of DHPS in bacterial metabolism and its absence in humans makes it a preferred antibacterial target. However, microorganisms like *Staphylococcus aureus* develop resistance to these antibiotics^18–20^ through mutations in the DHPS gene that lead to structural changes, reducing the enzyme’s affinity for sulfonamides and allowing the bacteria to continue folate synthesis despite the presence of the antibiotics. Notably, this resistance is particularly concerning in Methicillin-resistant S. aureus (MRSA), a pathogen responsible for severe community-acquired and nosocomial infections^21^. Here, we investigate the evolutionary dynamics of Staphylococcus aureus Dihydropteroate Synthase (SaDHPS). We aim to identify DHPS substructures that contribute to DHPS evolvability and result in anti-folate resistant bacterial phenotypes. Previous studies have explored both pABA and pterin binding sites on DHPS as promising drug target regions for novel inhibitors/small molecules^22^. Biophysical studies have shown that DHPS mutations, particularly in residues interacting with pABA, often create steric hindrance for sulfonamide binding while maintaining pABA interactions, allowing continued folate synthesis^19^. These mutations enable DHPS to evade antibiotic selection pressure and develop resistance to antibiotics. Consequently, traditional drug development efforts often fail to address the challenges posed by rapid mutations, which significantly contribute to the emergence of antibiotic resistance. Therefore, a comprehensive understanding of how DHPS adapts to selection pressure is critical for identifying novel vulnerabilities and developing effective therapeutic strategies.

This study proposes an *in-silico* strategy to identify the region which presented the highest evolutionary as well as structural impact on S. aureus DHPS (SaDHPS). Any disruption in this region would compromise structural stability, catalytic activity, and the resistance mechanism. The evolutionary investigation revealed highly interdependent residue positions whereas structural analyses provided their correlated motions/fluctuations, pairwise interaction patterns as well as how mutations influence their interactions with other residues within the protein structure. Our network-based analysis identified clusters of highly correlated residues, further elucidating their interaction network and structural significance. We performed Pareto Optimality analysis to understand the significance of the evolution and structure-based properties towards the structural dynamics of the protein. Further, we integrated those evolutionary features with the structural properties by Principal Component Analysis (PCA). We identified the Structural Blocks (SB3) that contains higher evolutionary and structural features. This region would be critical in regulating the resistance mechanism of the protein variants towards antibiotics. It makes SB3 a plausible antibiotic target region which effectively would disrupt/hinder the resistance possibility. The findings presented in this study contribute to ongoing efforts to develop drugs that can effectively counteract resistance mechanisms.

## Results

In this study, we employed a comprehensive approach that integrates both evolutionary and structural features to identify a sub-structure within the Staphylococcus aureus Dihydropteroate Synthase (SaDHPS) that plays a pivotal role in regulating the protein’s evolvability. Our objective was to identify a region within the protein that, if targeted, could disrupt the mechanism by which resistance to antibiotics emerges. This integrated strategy allowed us to map out evolutionary constraints and structural dynamics that collectively govern the stability and function of SaDHPS, thereby identifying a region that could serve as a promising target for therapeutic intervention. The research workflow, depicted in Fig. 1, outlines the *in-silico* strategy applied to SaDHPS, a bacterial model protein system. We performed extensive sequence space analysis of SaDHPS, to identify evolutionarily interdependent residue positions. These positions are critical, as they form evolutionarily selected pair-wise residue contacts that play key roles in maintaining the structural integrity and function of the protein under selective pressures. By mapping conserved and interdependent residues, we gained insights into the evolutionary pressures acting on SaDHPS and identified positions that are less tolerant to mutations, making them potential weak points in the protein’s defense against antibiotics.

**Figure 1:**
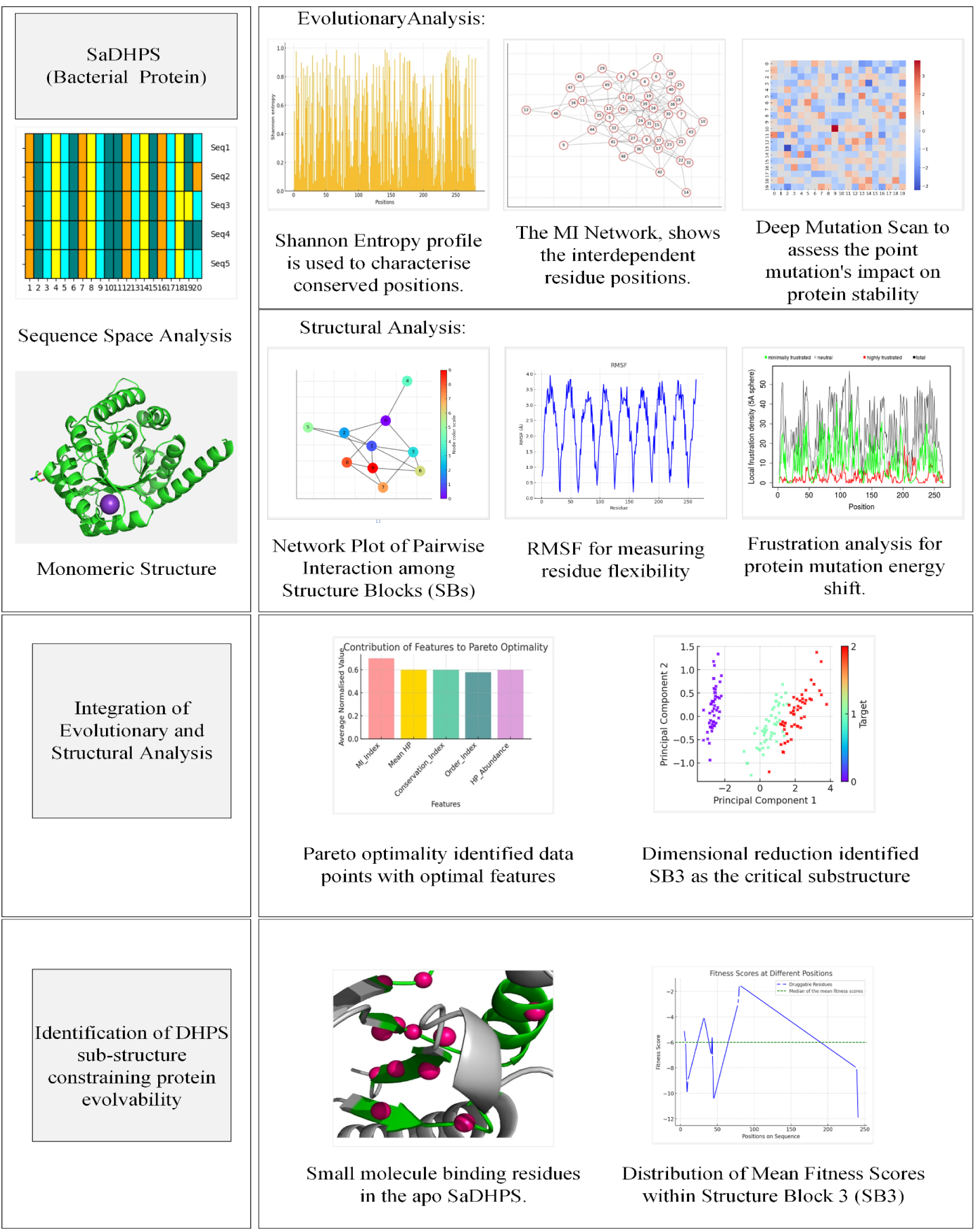
Workflow for identifying evolutionary and structural constraints in Staphylococcus aureus Dihydropteroate Synthase (SaDHPS). The sequence space analysis involved identifying conserved residue positions using Shannon Entropy profiling, constructing a Mutual Information (MI) network to reveal interdependent residues, and performing a deep mutation scan to assess the impact of point mutations on protein stability. Structural analysis included network plotting of pairwise interactions among structure blocks (SBs), RMSF analysis for residue flexibility, and frustration analysis to evaluate energy shifts caused by mutations. Integration of evolutionary and structural data through Pareto optimality analysis and Principal Component Analysis (PCA) identified Structure Block 3 (SB3) as a critical sub-structure with significant constraints. Druggability analysis of SB3 revealed small molecule binding residues, with a distribution of mean fitness scores indicating regions that are evolutionarily constrained and crucial for limiting protein evolvability, thereby providing a potential target for disrupting antibiotic resistance.

After the sequence space analysis, we delved into the mutational landscape of SaDHPS, focusing on how point mutations affect these evolutionarily co-varying residue positions. By examining the mutational landscape, we predict the impact of various mutations on protein function and stability, highlighting regions within the protein that are particularly sensitive to changes. These analyses provided a deeper understanding of the mutational tolerances of SaDHPS and identified regions where mutational changes could impact the protein’s ability to evolve resistance. Our structural analysis phase involved a detailed exploration of the protein’s structure network, Monte Carlo (MC) simulations, and frustration calculations. The structure network analysis enabled us to map the interactions between residues, while MC simulations provided insights into the dynamic behavior of the protein. Frustration calculations further elucidated areas within the protein where conflicting interactions could lead to structural instability. Together, these structural analyses allowed us to understand how the evolutionarily interdependent residues influence the overall dynamics and interaction patterns within SaDHPS. This information was critical in identifying the sub-structure that acts as a regulatory node, constraining the protein’s ability to evolve in response to selective pressures.

To integrate the evolutionary and structural data, we employed Pareto optimality analysis and Principal Component Analysis (PCA). Pareto optimality analysis identified regions of the protein that represent a trade-off between evolutionary conservation and structural flexibility. PCA allowed us to identify a specific sub-structure within SaDHPS that is crucial for maintaining both its structural integrity and its resistance to evolutionary changes. This sub-structure emerged as a key regulatory element in the protein’s ability to adapt, making it a prime target for interventions aimed at disrupting the emergence of antibiotic resistance.

Finally, we conducted druggability analyses to assess the therapeutic potential of the identified sub-structure. This analysis involved evaluating the physical and chemical properties of the sub-structure to determine its suitability as a binding site for small molecule inhibitors. The results of this analysis suggest that the identified sub-structure possesses the necessary characteristics to be a viable drug target.

### Sequence space analysis has provided evolutionarily critical positions and mutational tolerance

Throughout the evolutionary timescale, the residues that do not change or alter very slowly are conserved residues. In order to understand the conservation propensity of amino acid positions in the DHPS sequence, we performed sequence conservation analysis and quantified using Shannon’s entropy (SE) calculation. In the SE profile, an increase in spike heights, i.e., higher SE values, represents lower positional conservation. The SE profile of DHPS shows that evolutionarily conserved positions (with low or zero SE score) are scattered throughout the protein sequence (Fig. 2A).

**Figure 2:**
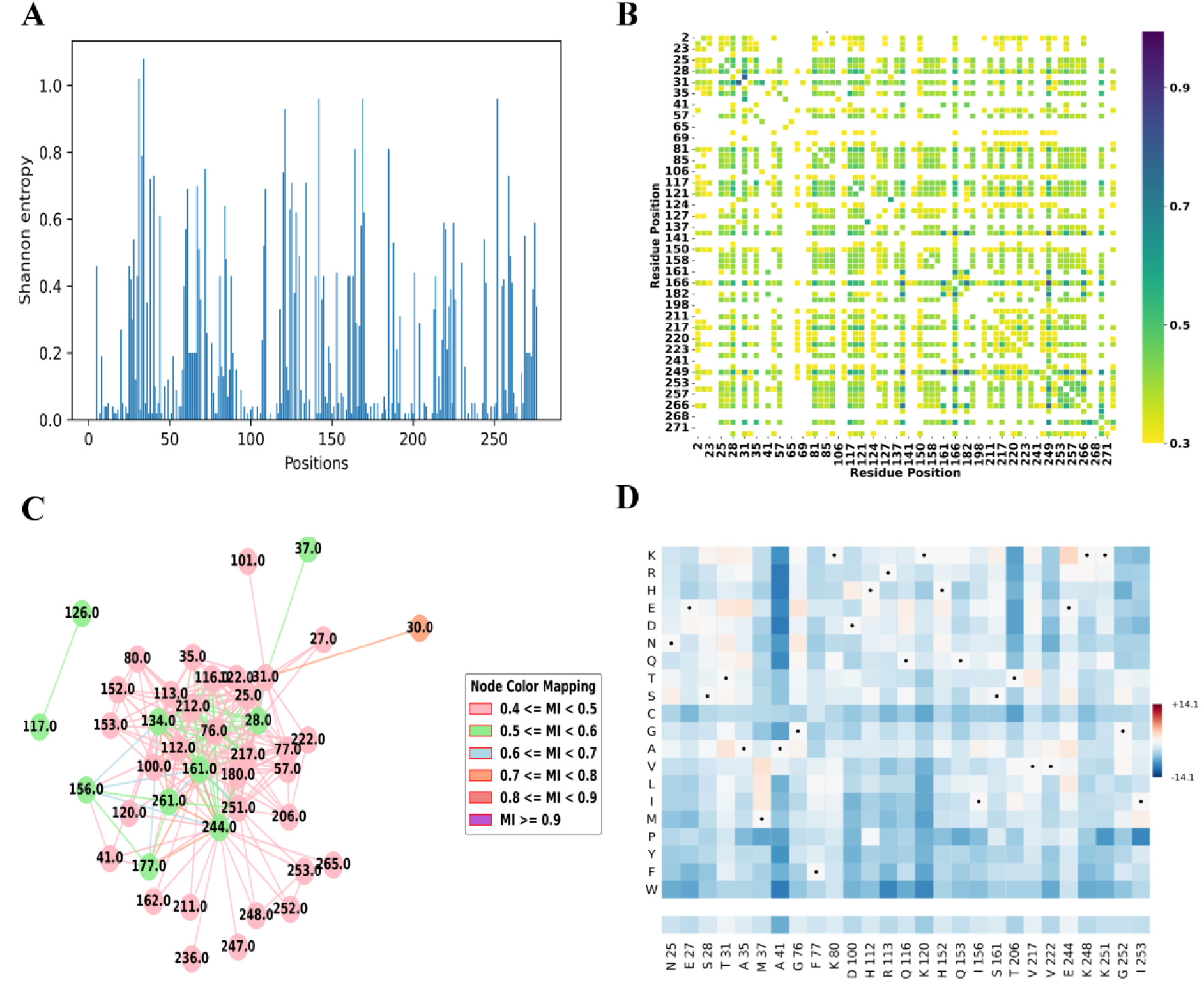
Sequence space study of DHPS identifies important regions, which are under evolutionary restraints. **(A)** Shannon Entropy profile showing positional conservation pattern. High SE values (Spike heights) correspond to lower conserved residue positions and vice-versa. **(B)** Correlation matrix heat-map showing mutual information values obtained from multiple sequence alignments. **(C)**Mutual Information network (MI Network) showing the interdependent residue positions. A dense connection indicates higher interdependence of that residue position on others. (D) Phenotypic effects of point substitutions on the 28 interdependent residue positions. Each of the residue positions is substituted with the other 19 naturally occurring amino acids individually and the mutational impact was calculated, forming a row in the heatmap. The identity of each row is given by the single letter amino-acid code given on the left of the heatmap. The intensity of the blue color is proportional to the damaging effect on fitness, whereas the shades of white show neutral effects, and dark red indicates increasing beneficial effects on the fitness of the protein. The amino acid originally present in the DHPS sequence at each position is demarcated by a black dot for that corresponding amino acid row in the heatmap.

However, mutations can happen in the non-conserved regions of the protein. If two residues of a protein are in proximity in the 3D structure, the effect of destabilizing amino acid substitution at one position would be nullified by a mutation of the other position over the evolutionary timescale in order to maintain the interaction between the residue pair^23^. Such compensation would determine positional interdependence in a protein sequence, which are extremely critical in defining the constraints in the evolutionary sequence space. We identified these evolutionary interdependent positions by performing Mutual Information calculation (MI calculation). High MI value for a co-evolving pair indicates higher positional interdependency, indicating their strong interaction towards contact formation. On analyzing the MI profile, we considered the top 50% of the interdependent residue pairs (MI pairs) having MI scores from 0.43 to 0.8 (Fig. 2C). We obtained a total of 28 highly co-varying residue pairs throughout the sequence of DHPS, which impose contacts in the 3D structure (Table 1) (Fig. 2B and 2C). Fig. 2C illustrates the intricate network of 28 highly interdependent residues within the DHPS sequence. The dense connectivity observed in the network highlights the significant extent of interdependency among these residues. We employed different color codes to represent varying ranges of interdependency values, facilitating a clear visualization of the inter-residue relationships. The color gradient allows for the differentiation between low, moderate, and high inter-dependencies, thereby providing insights into the evolutionary co-varying residues and their influence on the structural coherence of the protein.

**Table 1:**
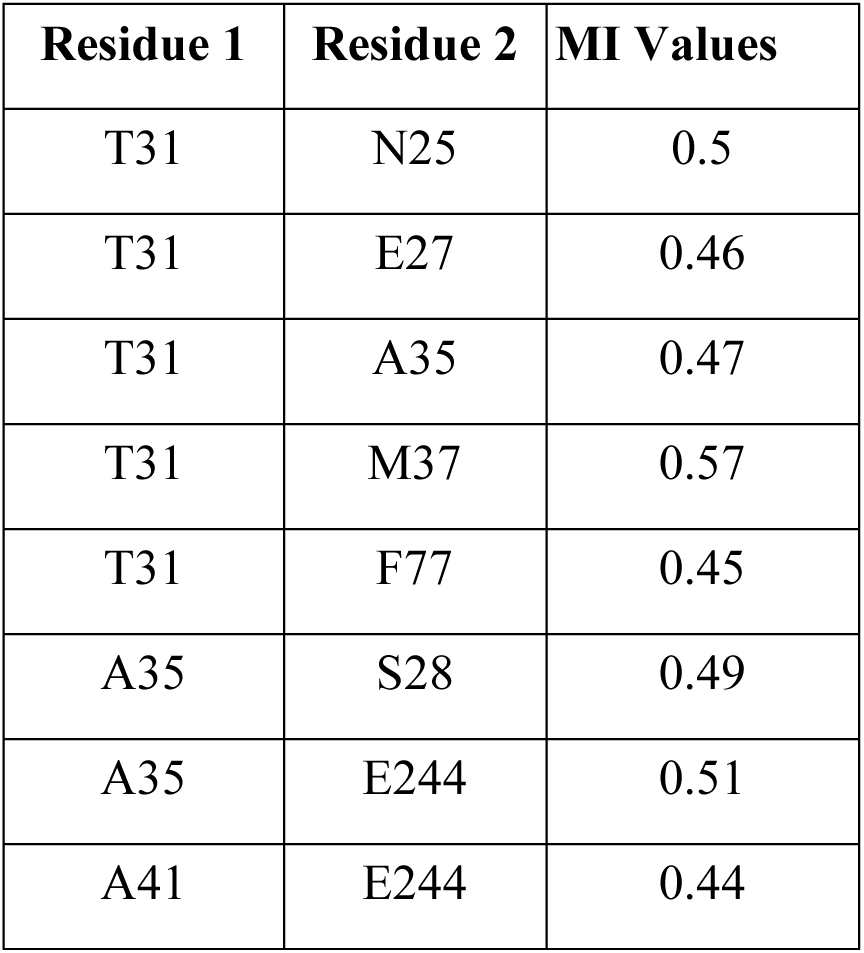
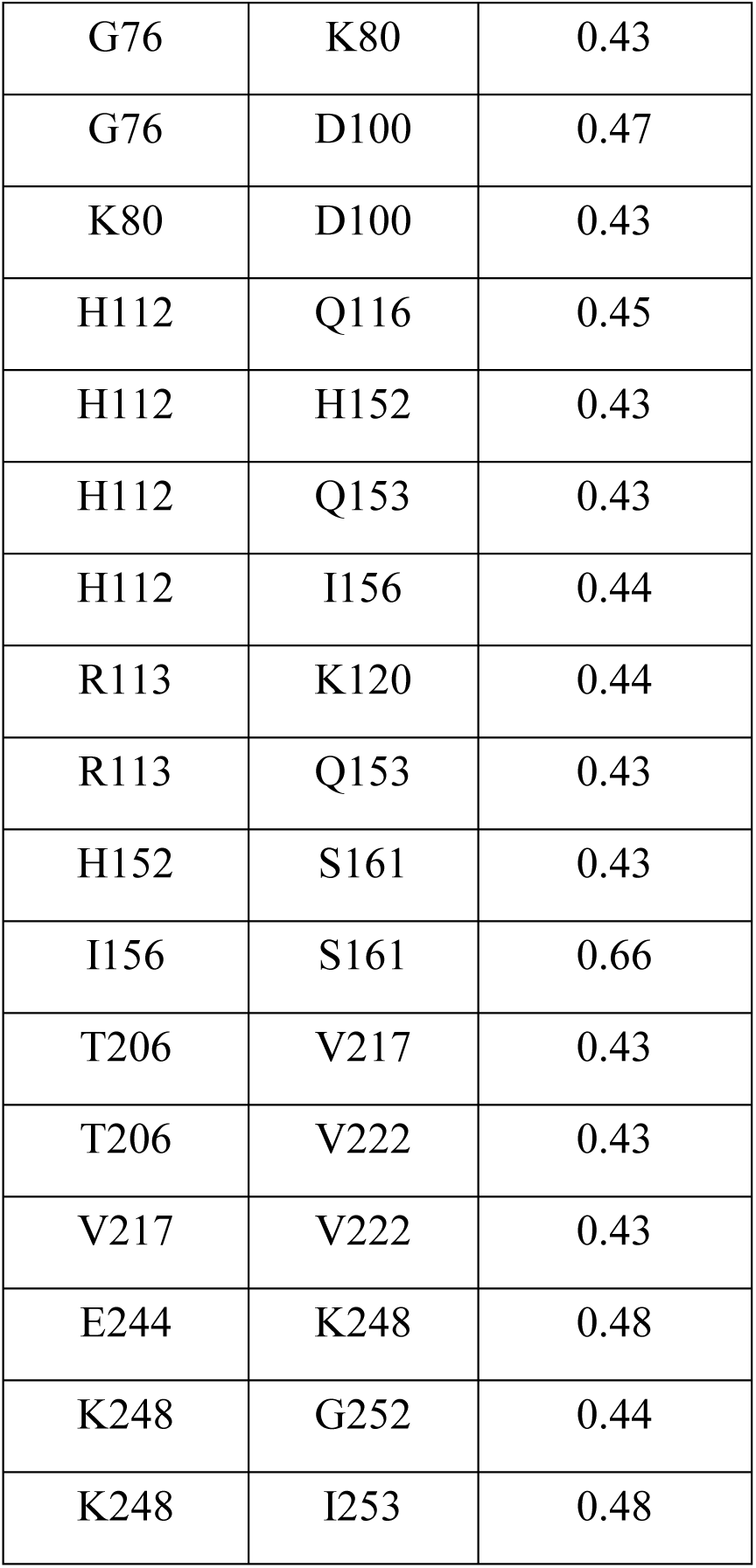
Top MI pairs

| Residue 1 | Residue 2 | MI Values |
| --- | --- | --- |
| T31 | N25 | 0.5 |
| T31 | E27 | 0.46 |
| T31 | A35 | 0.47 |
| T31 | M37 | 0.57 |
| T31 | F77 | 0.45 |
| A35 | S28 | 0.49 |
| A35 | E244 | 0.51 |
| A41 | E244 | 0.44 |
| G76 | K80 | 0.43 |
| G76 | D100 | 0.47 |
| K80 | D100 | 0.43 |
| H112 | Q116 | 0.45 |
| H112 | H152 | 0.43 |
| H112 | Q153 | 0.43 |
| H112 | I156 | 0.44 |
| R113 | K120 | 0.44 |
| R113 | Q153 | 0.43 |
| H152 | S161 | 0.43 |
| I156 | S161 | 0.66 |
| T206 | V217 | 0.43 |
| T206 | V222 | 0.43 |
| V217 | V222 | 0.43 |
| E244 | K248 | 0.48 |
| K248 | G252 | 0.44 |
| K248 | I253 | 0.48 |

**Table 2.**
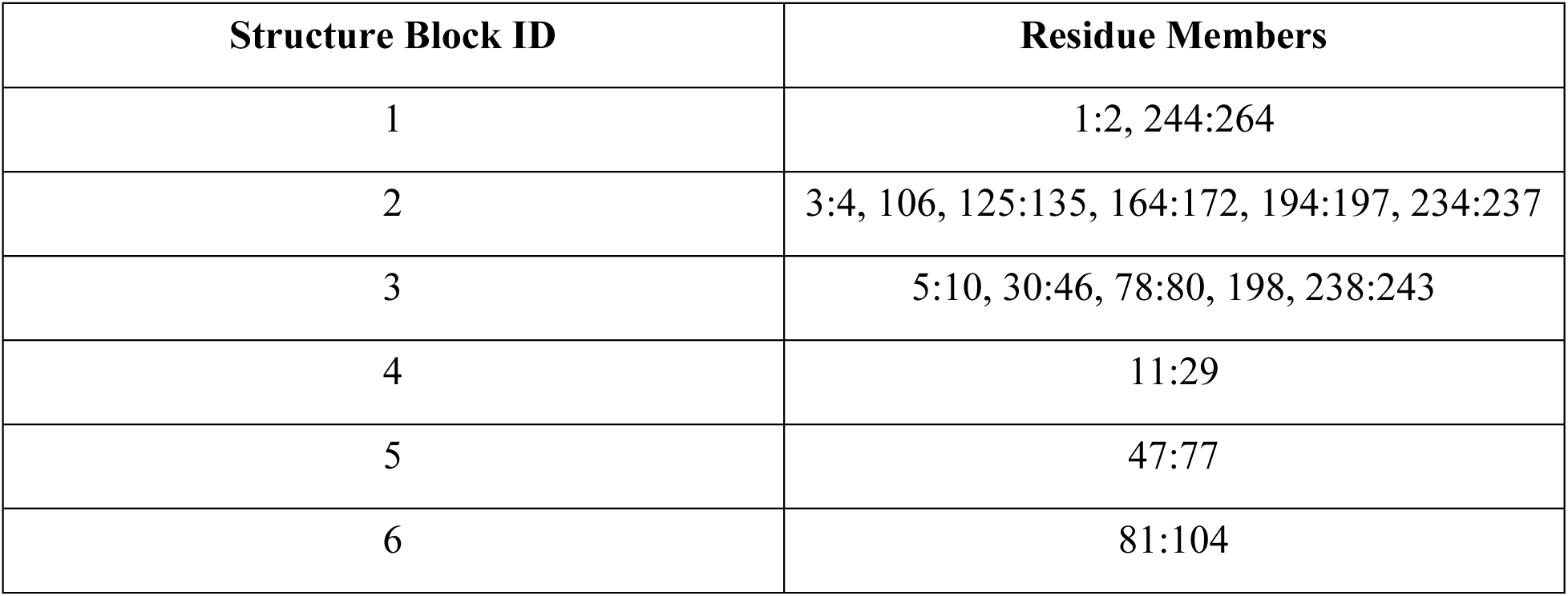
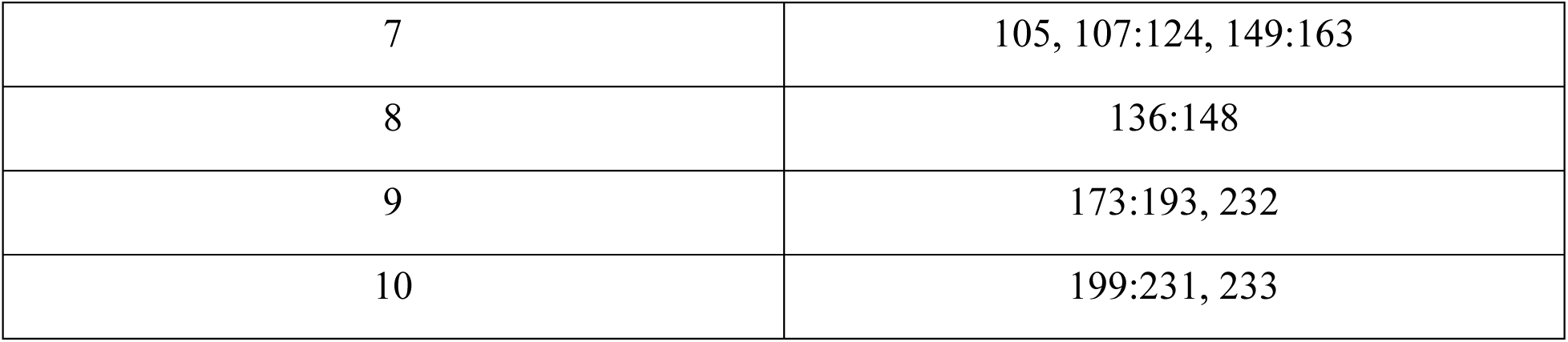
DHPS Structure Block Members

From the MI profile, we found six interdependent residue positions (25, 27, 28, 31, 35, and 37) (Table 1) are in the helix which is adjacent to loop I. The mobility of loop I (residues 11 - 25) and loop II (residues 47 - 59) are crucial for the catalysis of the enzyme by helping in the para amino benzoic acid (pABA) binding pocket formation^19^. Interestingly, we observed evolutionary interdependent residue positions at the dimer interface of the protein, with four (212, 222, 244, and 251) of them forming contacts across the monomers and, in turn, modulating their movements. Also, we found that residue position 206 has higher interdependency with 217 and 222. These residues are in close proximity to residue 203, which can interact with both the substrates (DHPt and pABA). Therefore, the evolutionary analyses clearly indicated that evolutionarily co-varying residues play a crucial role in dimerization of the protein, the movement of loop I and II as well as in substrate binding. These findings highlight the structural as well as functional importance of evolutionarily co-varying residues and their potential impact on the protein’s biological activity.

Apart from providing us with the evolutionary history of a protein, the sequence also encrypts information regarding the impact of mutations on the protein structure and its overall stability. In order to assess the possible effect of a point mutation on the overall stability of the protein structure, we performed Deep Mutation Scan and generated a mutational landscape. This landscape provides us with information on positional tolerance and the potential effect of substitution on the biophysical fitness of the protein structure. We observed that 50 % of the interdependent positions had near-neutral effects (towards white) for multiple substitutions of different amino acid types (25, 28, 31, 76, 77, 80, 112, 152, 161, 206, 217, 244, 251 and 252) (Fig. 2D). We also found that seven residue positions (25 % of the interdependent positions) having more than one substitution indicated to be beneficial (darker shades of red) in terms of protein stability (31, 35, 37, 76, 116, 152, and 244). These observations show that the interdependent residue positions are very tolerant towards accommodating point mutations. Additionally, a significant proportion of the interdependent positions may have a prominent role in enhancing the stability of the protein and the corresponding fitness of the organism (25% of the interdependent positions). Mutations in these residue positions are usually followed by a compensatory mutation in the other position of the co-varying pair to maintain the interactions between them.

### Integrated Structural Analysis to identify DHPS substructure critical to stability

By performing sequence space analysis, we identified evolutionary interdependent positions in DHPS. Next, we analyzed the structural details of DHPS to reveal the important region that would modulate the internal dynamics of the protein.

To understand the internal orchestration of the protein in terms of their correlated motions, we performed a structure network analysis^24^. We generated all residue networks of the apo structure of SaDPHS and split it into a simplified community cluster network by applying the Girvan-Newman clustering algorithm. We found that the protein structure comprised 10 structure blocks (SBs) (Fig. 3A).

**Figure: 3.**
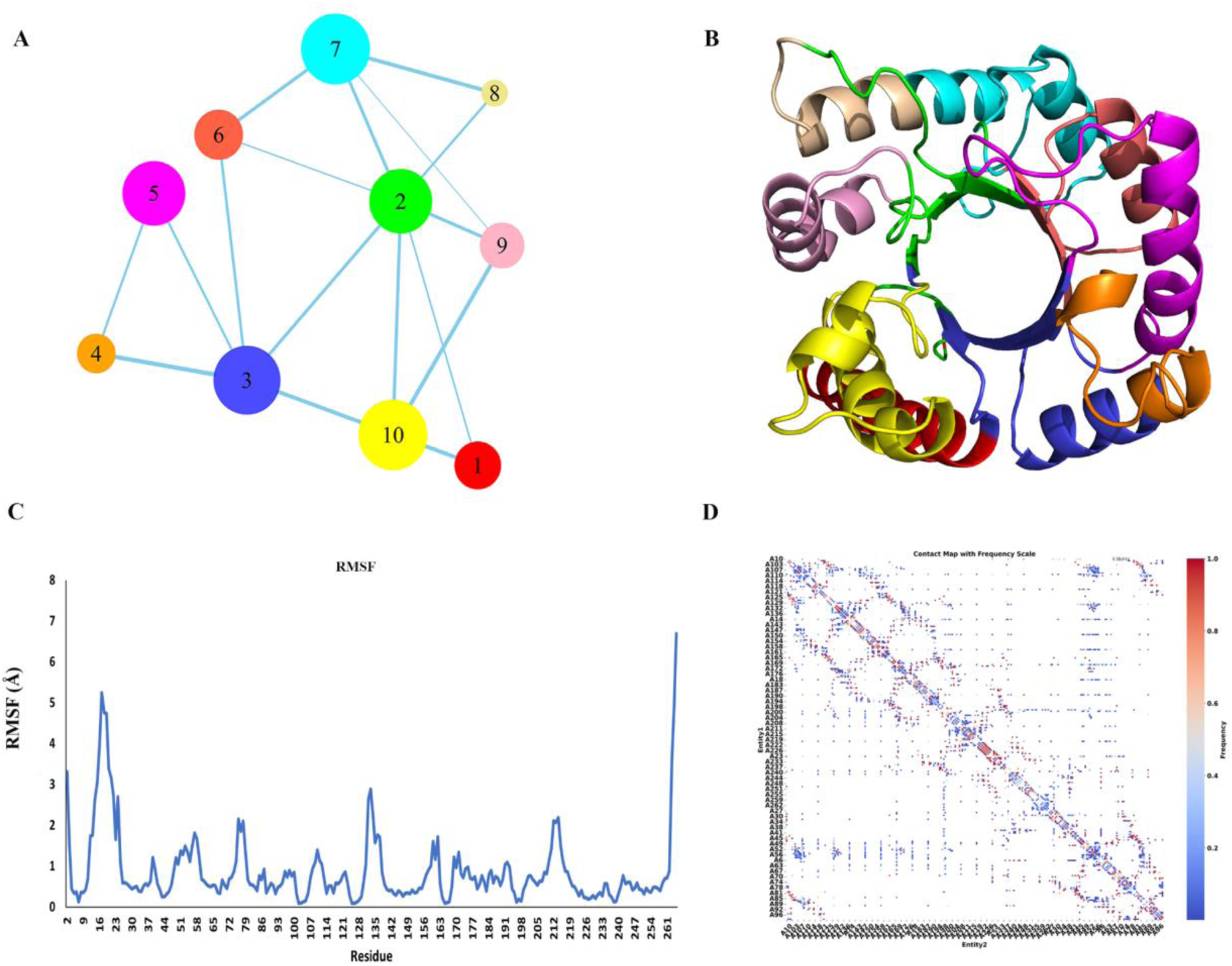
Structural investigation of DHPS to identify the region that plays a critical role in regulating loop dynamics. **(A)** Structure Network plot of SaDHPS. In the community-cluster network, the nodes represent different structure blocks (SBs) that are comprised of groups of residues, and edges are the correlation of the residue motions. The width of the edges dictates the extent of interaction between the SBs. The SaDHPS structure has been found to be comprised of 10 SBs. **(B)** Crystal Structure of the apo form of SaDHPS (PDB I’D: 1AD1). The colors represent the residues of the corresponding Sbs. **(C)** RMSF values calculated from NMA. These corroborate the high mobility of the loop 1 region (residues 11-25). (**D**) Contact map depicting the interaction between residues within all protein structures. Residue numbers are shown on the axes, and each point of the map indicates a pair of residues that are in close proximity (within a cutoff distance) during the simulation. More frequent or probably higher encounters are indicated by darker hues. This map provides insights into the flexibility of the protein by highlighting critical interactions that are necessary for both its structural integrity and function.

Analyzing these 10 SBs of SaDHPS, we observed that loop I and loop VII residues were housed in SB 4 and 5, respectively. The motion of these two loops (Fig. 3C, D) leads to the transient formation of the pABA binding site^19^. Interestingly, they are separated from the rest of the proteins in the community cluster network. The structure network shows that SB 3 acts as a bridging cluster, connecting loop I and loop VII (SB 4, 5) to exchange allosteric impact between them and the rest part of the enzyme, particularly with SB 2 housing the DHPt-PP binding residues (Fig. 3A). In addition, we found that the Mn^2+^ ion-binding residue, i.e., Asn11^15^ is located in SB 4. Also, we observed that the residues at the dimer interface were in SB 1 and 10. Apart from SB 3, SB 2 has the highest number of connections in the network, followed closely by SB 7. Here, we considered/selected/chose seven SBs (1, 2, 3, 4, 5, 7, and 10) that have a higher number of connections in the network as well as the residues of these SBs are important in terms of the catalysis of DHPS.

We also looked at the distribution of the mean disorder propensity (MDP) calculated using RIDAO^25^ within the amino acid sequence of SaDHPS (Uniprot ID - O05701). Results of this analysis are shown in Fig. 4, which compares the distributions of RMSF, Shannon entropy, and MDP score over the SaDHPS sequence and also represents positions of the 10 SBs. Analysis of this figure shows that although SaDHPS is predicted as mostly ordered protein (most of its residues have MDP scores below 0.5), structure of this bacterial enzyme is expected to be rather flexible, as its sequence averaged MDP score is 0.27±0.14. Therefore, SaDHPS is classified as an ordered protein based on the accepted classifications of proteins based on their percent of predicted intrinsically disordered residues (PPIDR), where proteins are considered as ordered, moderately disordered, or highly disordered, if their PPIDR < 10%, 10% ≤ PPIDR < 30%, or PPIDR ≥ 30%. However, according to the Average Disorder Score (ADS)-based grouping, where ADS < 0.15, 0.15 ≤ ADS < 0.5, and ADS ≥ 0.5 values are used to classify proteins as ordered, moderately disordered/flexible, and disordered, respectively, this enzyme is classified as moderately disordered. Fig. 4 shows that in addition to disordered N- and C-termini, SaDHPS contains three flexible regions with 0.15 ≤ ADS < 0.5. These flexible regions act as “envelopes” that include regions with higher Shannon entropy and higher RMSF. The only noticeable exception is given by the most ordered region of the protein (residues 80-120) that includes most of the SB 6, and parts of SB 2 and SB 7, and where distributions of MDP, Shannon entropy, and RMSF are not correlated. Fig. 4 also shows generally good correlation between the Shannon entropy and RMSF, as typically regions with high RMSF are characterized by higher evolvability or at least located in the close proximity to the regions with higher Shannon entropy. Iinteresting examples are given by the low RMSF regions 25-36, 57-65, 177-184, 221-227, and 243-254, which contain residues with high Shannon entropy. There are also three short regions with high RMSF and low Shannon entropy (residue 18-21, 105-109, and 163-176). These observations indicated that as a rule, highly evolvable residues are located within flexible regions characterized by higher RMSF values and/or higher MDP scores.

**Figure 4.**
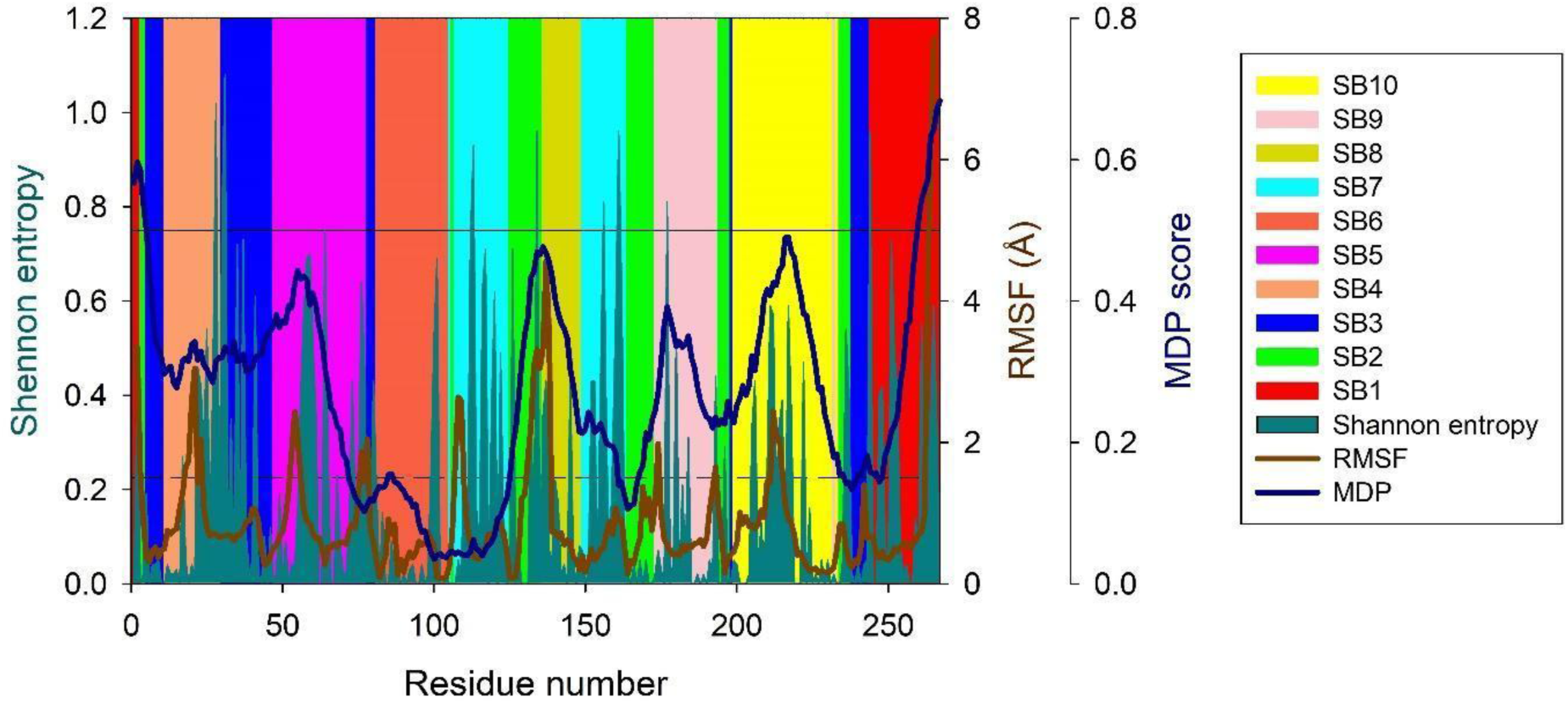
Comparison of distributions of MDP, RMSF, and Shannon entropy within the SaDHPS sequence. Positions of 10 SBs are shown by differently colored areas.

In order to quantify the localized frustration within the SaDHPS structure, we calculated the frustration generated due to interactions among the residues and classified them as minimally, neutrally, or highly frustrated. We observed that SB 2 had a total of 251 interactions, out of which 7.17% (18 interactions) generate higher extent of frustration to the overall structure of the protein (shown as red color) (Fig. 5A). Also, 39.84% of the inter-residue interactions (100 interactions) create minimal frustration (green) that contribute to the stability of the protein. SB 3 residues consisted of a total of 286 interactions with other residues (Fig. 5B). Interestingly, around 48.6% (139) of these contact produced minimal frustration. 125 amino acid contacts (43.71%) generated neutral frustration, while only 22 contacts (7.69%) were responsible for high frustration in the structure. The SB 3 residues interacted well with other significant regions of the protein. It showed that mostly the inter-residue interactions generating minimal or neutral frustration involved Loop I (11, 12), Loop II (47, 48), and dimer interface residues (244, 251) (Fig. 5B). In case of SB 7, out of 215 interactions, 8.84% (19 contacts) generated high frustration, 50.7% and 40.47% of the interactions created neutral and minimal frustration. Notably, none of the SB 7 residues interacted with other areas of protein, whereas SB 2 residues had few contacts with the Dimer interface, the majority of which were highly frustrated, and SB 3 had the most favorable interactions with minimal and neutral frustration states.

**Figure 5.**
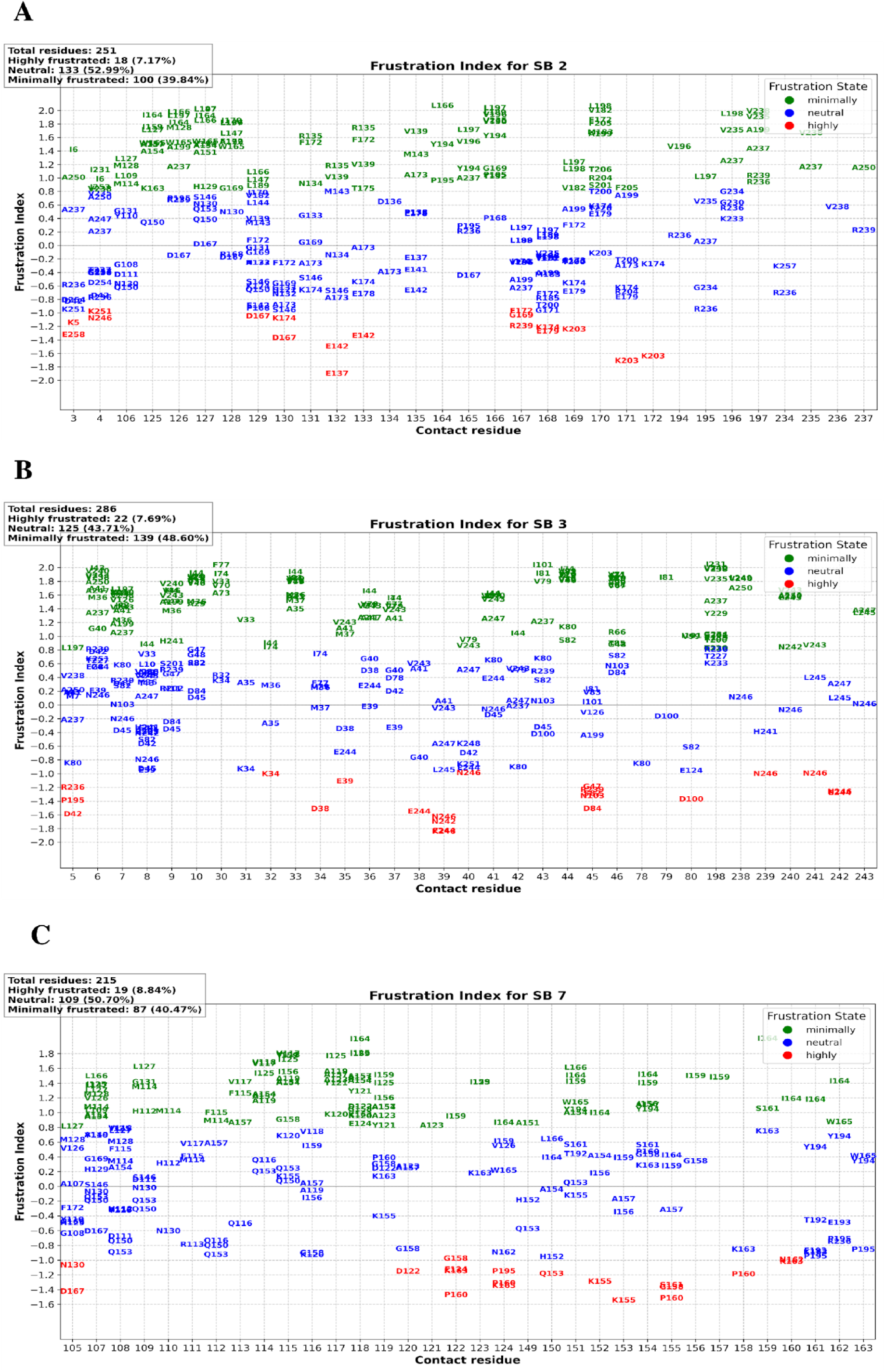
The frustration index values of residue contacts based on protein mutational energy changes are shown in this plot. (**A**) Structural Block 2 frustration index for all 251 mutational changes. (**B**) Structural Block 3 frustration index for all 286 mutational changes. **(C)** Structural Block 7 frustration index for all 215 mutational changes. Matching frustration index values are shown on the y-axis, while residue pairings are shown on the x-axis. Each point represents the favorability of a contact; higher values denote more favorable connections.

### Combined Sequence and Structure Space Information to Identify a Critical Region in DHPS

From our sequence space analysis of SaDHPS, we observed 28 interdependent residue positions having higher mutational tolerance. From the structural analysis, we identified seven SBs that either have key residue positions important for the catalysis or have higher numbers of inter-residue connections. Here, we delved into the ensemble of evolutionary and structural features within our dataset through Pareto optimality analysis along with the Principal Component Analysis (PCA). We aimed to identify the structure block that represented optimal combinations of these features, where no single index or features could be improved without compromising another. Here, we identified five Pareto optimal points, highlighting structure blocks characterized by a distinct balance of indices.

The Pareto Optimality analysis revealed that MI Index, with a 25.08% contribution, emerged as the most influential feature in determining Pareto optimality, followed closely by Mean HP and Conservation Index (Fig. 6A). This finding clearly indicates the significance of Mutual Information Index in the optimality landscape of the DHPS protein. In terms of Structure Blocks (SBs), our utility model highlighted SB 3 as the most important, having the highest utility score of 0.8076 (Fig. 6B). This SB displayed an optimal combination of the identified critical features, particularly excelling in MI Index. Following SB 3, SBs 1, 5, 10, and 7 were also identified as significant, each demonstrating a strong alignment with the critical features but with varying proportions (Fig. 6B). These SBs represent scenarios in the dataset where the harmonization of key features culminates in a state of Pareto optimality, serving as benchmarks for other data points (Fig. 6C). Additionally, the ’Distribution of Utility Scores’ plot offered a broader perspective, displaying the range and frequency of utility scores across the dataset, thereby contextualizing the exceptional nature of the top-scoring SBs (Fig. 6D).

**Figure 6:**
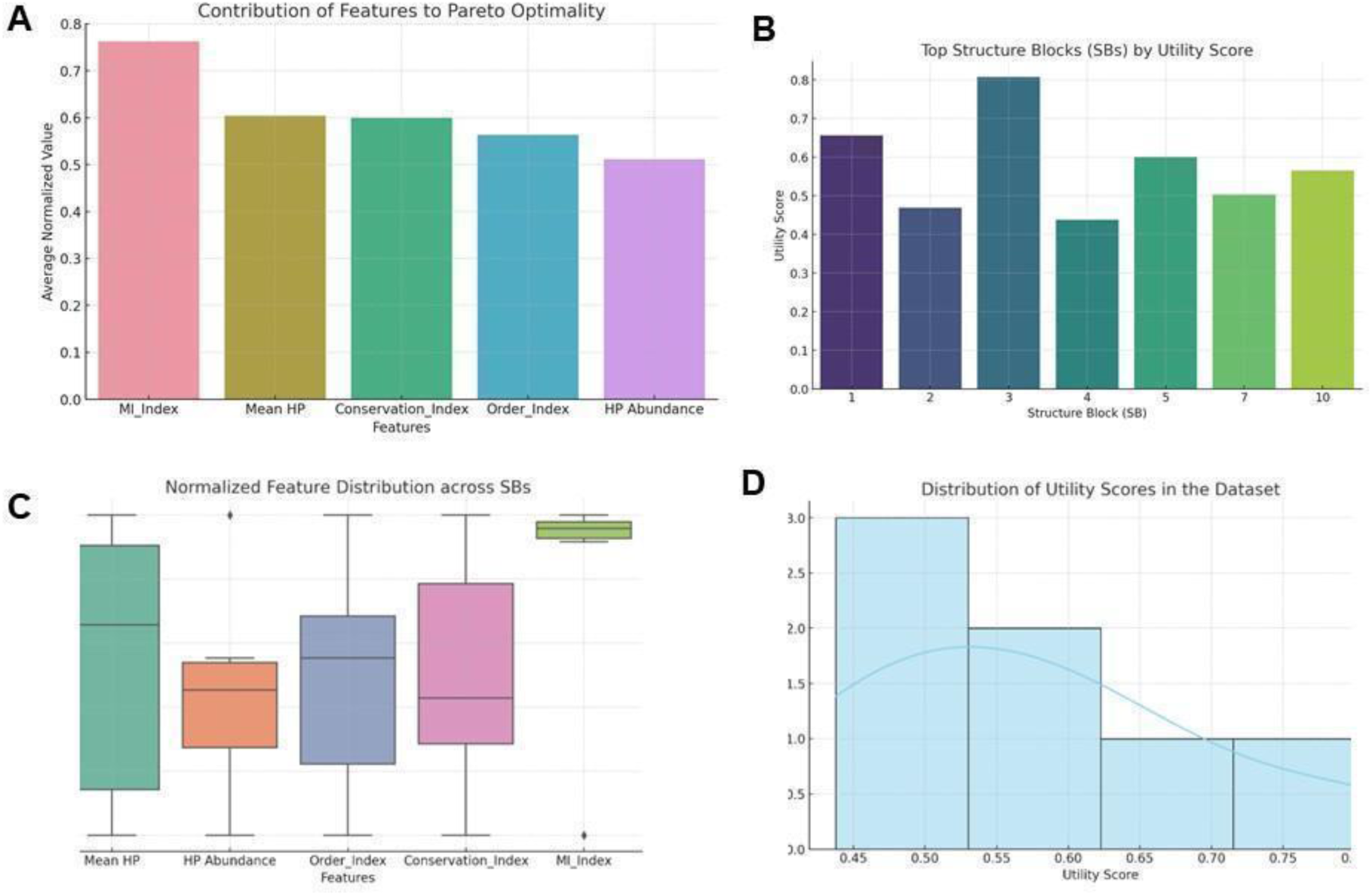
Pareto Optimality analysis unveils the pivotal role of features in determining optimality within the DHPS protein landscape. (A) the Mutual Information (MI) Index contributes 25.08% to Pareto optimality, closely trailed by Mean HP and Conservation Index. (B) Structure Block 3 (SB 3) displays a utility score of 0.8076. SBs 1, 5, 10, and 7 also demonstrate significance, each displaying varying alignments with critical features. (C) Normalized Pareto optimality feature distributions among different SBs shows how the key features leads to an optimal state within the dataset. (D) ’Distribution of Utility Scores’ plot shows the range and frequency of utility scores across the dataset.

In order to perform PCA, we integrated information from the sequence space and structural analysis in a common, hybrid subspace. This also would ensure that we captured the essential sequence and structural features of each SB and would validate our Pareto optimality analysis. Here, we considered four sequence and structure based indices (see methods) and applied dimensionality reduction by means of PCA. PCA linearly projects high dimensional data to a low dimension space having a few, uncorrelated principal components. We reduced the sequence and structure based indices individually and visualized the shortlisted SBs (SB1, 2, 3, 4, 5, 7 and 10) (see methods) on this sequence-structure hybrid subspace (Fig. 7A). The PCA plot shows that SB 3 and SB 7 have a wide overlap with most of the SBs across the sequence and structure components (Fig. 7A and 7B). This shows that both SB 3 and SB 7 have similar evolutionary and biophysical properties like the majority of SaDHPS.

**Figure 7:**
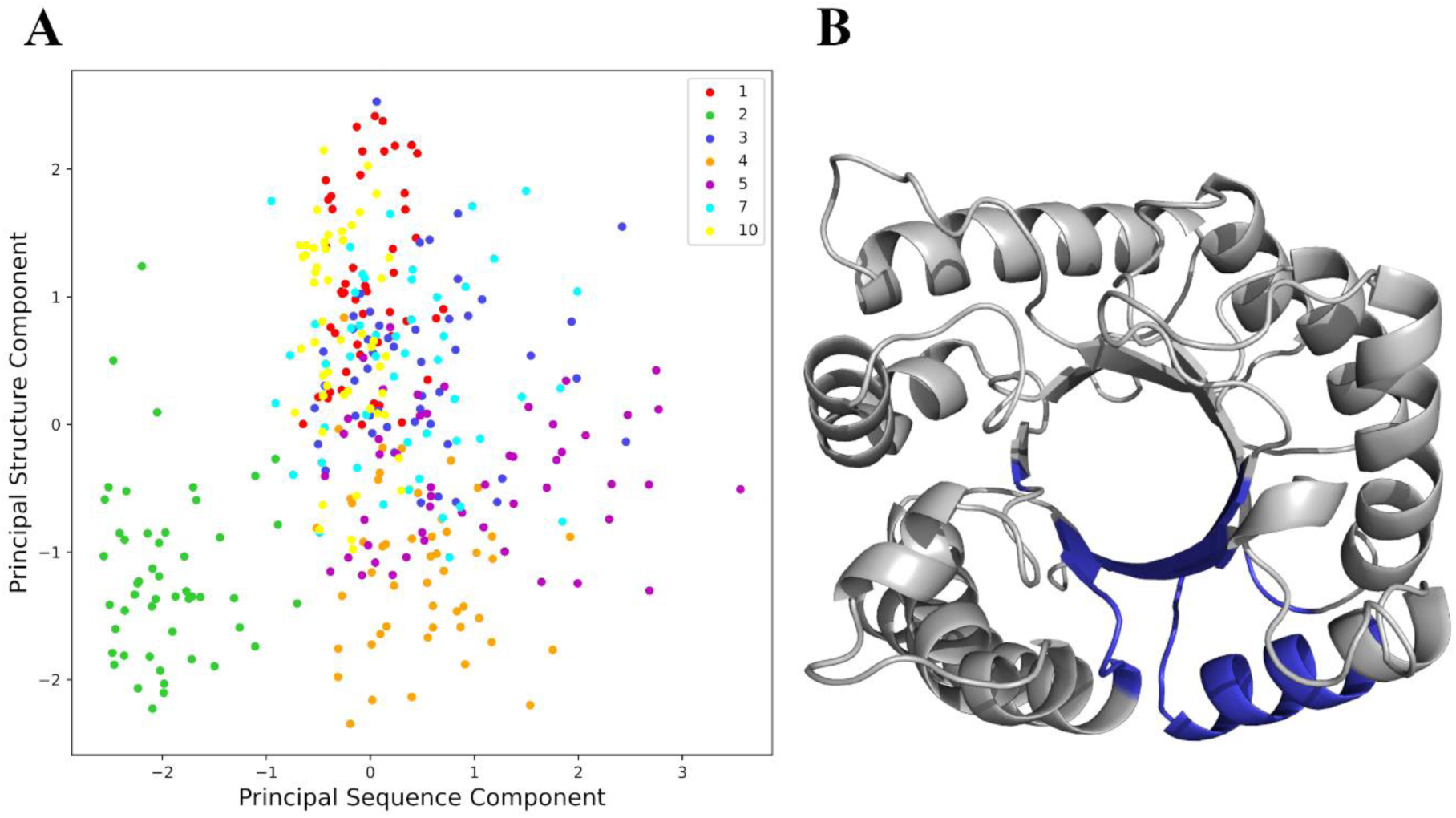
Principal Component Analysis (PCA) combining evolutionary as well as structural properties of DHPS identified Structure Block (SB) 3 as the most important region of DHPS. **(A)** PCA plot of the shortlisted SBs from the structural analysis in the hybrid sequence-structure subspace. This subspace was created by reducing the dimensions of sequence and structure based indices (see methods) separately to make the principal sequence component and principal structure component respectively. The legend is given within the plot in the top right **(B)** Crystal structure of apo form of SaDHPS (1AD1) showing the region of SB 3 (blue).

### Pocket Validation using Computational Screening

Having established the evolutionary and structural significance of SB 3 in SaDHPS, we then assessed the druggability of the residues in this region. We searched for the drug binding residues within SB 3 using a variety of computational methods and ranked the obtained pockets for biologically significant binding sites via different physico-chemical parameters (explained in methods section). Computational geometry methods yielded/indicated residue 9, 10, 32, 45, 239 and 241 when we ranked the identified pockets according to the solvent accessible volume. We sorted out residue 5, 7, 42, 43, 78, 79 and 80 according to a composite druggability score. Protein threading methods identified residue 9, 239 and 241 by ranking them based on a confidence score and 9, 45, 239 and 241 by implementing a z score. We mapped these residues on to the structure of SaDHPS, which represents 40% of the entire residues of SB3, highlighting the presence of small molecule binding residues (SMBR) in SB3 (Fig. 8A). To make sure targeting SB3 with a drug is evolution-resistant, we checked the mutational propensities of the SMBR. We found that all of the residues have a negative effect on the stability of SaDHPS when substituted, with the median of the distribution close to -7.6, which point towards a low propensity of point mutations (Fig. 8B). This establishes the fact that therapeutically targeting the region SB3 of SaDHPS would be both effective as well as evolution-resistant.

**Figure 8:**
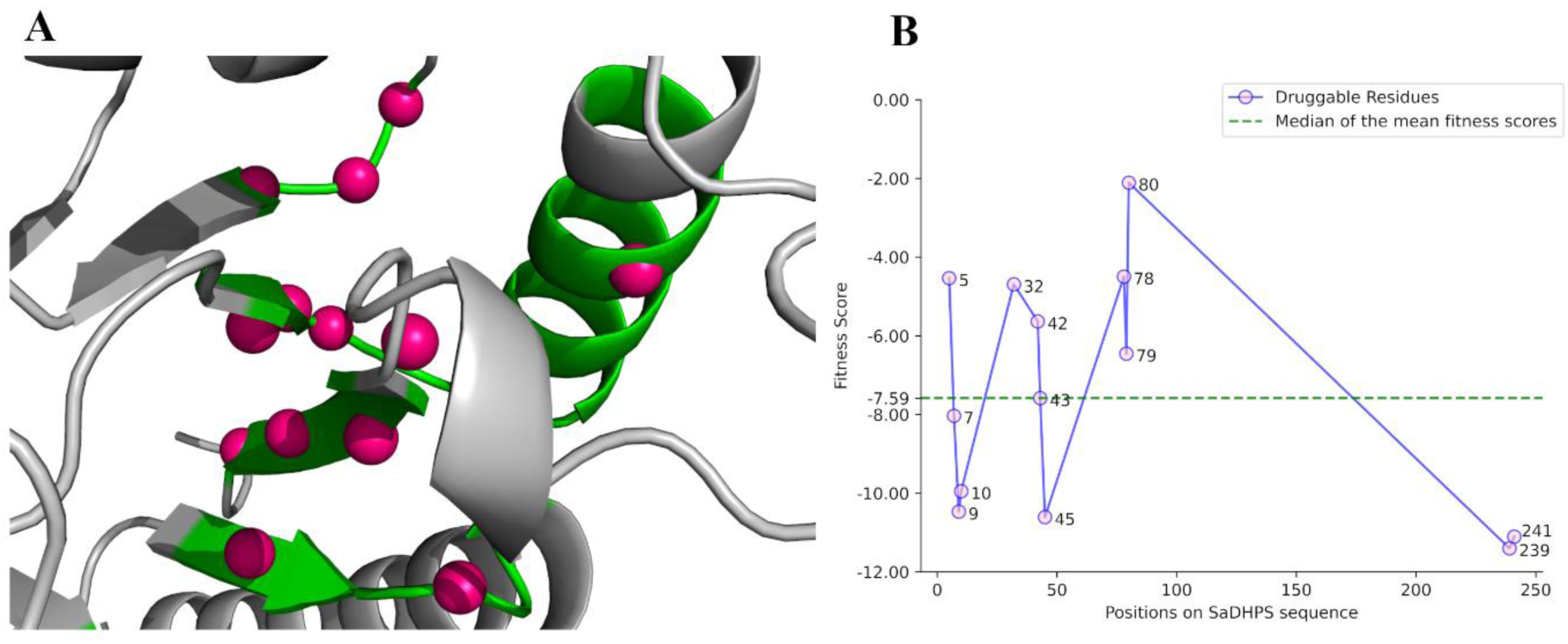
Identification of druggable residues in DHPS-structure-block-3 reveals low mutational tolerance among the constituent residues. **(A)** Small molecule binding residues (SMBR) shown as pink spheres within the apo form of SaDHPS (1AD1). The gray region indicates the rest part. Region encompassing SB3 is in green cartoon depiction. **(B)** Distribution of the mean fitness scores of the 13 SMBR within SB3, assessed by deep mutational scanning. A fitness score of zero represents no effect on the protein stability upon substitution. Negative values indicate a damaging effect while positive values indicate a beneficial effect. The median of the distribution (−7.59) is shown as a horizontal dashed line in the plot. The legend is given within the plot on the top right.

## Discussion

The rapid evolution of proteins under selection pressure is one of the primary causal factors for the emergence of antibiotic resistance, making it one of the most pressing challenges in modern medicine^20^. Pathogens, when exposed to antibiotics, undergo genetic changes that alter their protein structures, enabling them to survive and proliferate despite drug treatments^27^. This adaptive capacity, driven by the evolutionary principles of mutation, selection, and drift, emphasizes the need for a deep understanding of protein evolution to develop strategies that can effectively counteract resistance. Traditional drug development targets wild-type (WT) protein structures, making them ineffective against the escape mutants which evolve under drug induced selection pressure. However, emerging research emphasizes the importance of anticipating evolutionary trajectories, thus providing a proactive approach to drug development that could prevent the emergence of resistance^28^.

Analysis of protein evolvability can be an insightful marker for predicting protein and cell-systems level evolutionary dynamics^11^. Mutations enhancing protein evolvability not only improve the immediate fitness of an organism but also increase the likelihood that future mutations will be beneficial, thereby accelerating adaptation. This concept is particularly relevant in the context of antibiotic resistance, where rapid adaptation can render treatments ineffective. Previous work on protein and RNA fitness landscapes demonstrates how a small subset of mutations can significantly shift the adaptive potential of a protein, making subsequent mutations more likely to be advantageous^11^. Such insights suggest that targeting regions of proteins that are critical for their structural and functional integrity—yet less prone to accommodating beneficial mutations—could be a strategic way to inhibit the evolutionary flexibility that drives resistance.

In this study, we applied an integrated evolutionary and structural analysis to Dihydropteroate Synthase (DHPS), a key enzyme in the folate pathway and a well-established target of sulfonamide antibiotics. Our findings identified an allosteric, evolution-resistant sub-structure (or structure block) within DHPS, termed structure-block SB3 that plays a critical role in maintaining the enzyme’s structural stability and catalytic function. By combining sequence space analysis with structural dynamics, we demonstrated that this region is constrained in its ability to accommodate mutations, making it a prime target for therapeutic intervention. Targeting SB3 could destabilize the enzyme, reducing its capacity to evolve resistance to sulfonamides and offering a new avenue for antibiotic development. The identification of SB3 as a critical regulatory cluster within DHPS represents a significant advance in our understanding of how to combat antibiotic resistance. Unlike traditional approaches that focus on inhibiting the active site, we present a druggable pocket essential for the enzyme’s structure and function. Disrupting SB3 could prevent the formation of the pABA binding pocket, a crucial step in the enzyme’s catalytic cycle, thereby undermining the enzyme’s ability to function and adapt. This approach aligns with broader efforts to develop evolution-proof antibiotics by focusing on regions of proteins that are structurally constrained and less likely to tolerate adaptive mutations.

The implications of our findings extend beyond DHPS to the general understanding of protein evolution and antibiotic resistance. By targeting evolutionarily constrained regions, we introduce a new paradigm in drug design, where the focus shifts from merely inhibiting function to preemptively limiting the evolutionary potential of target proteins. This strategy not only addresses immediate resistance issues but also seeks to curtail the pathogen’s adaptive capabilities by exploiting inherent evolutionary vulnerabilities^29^. Such an approach is particularly important in the context of evolvability-enhancing mutations, which can rapidly shift the adaptive landscape and drive the emergence of resistance. The relationship between protein foldability and evolvability stresses the complexity of designing inhibitors that can withstand the evolutionary pressures exerted by pathogens. Proteins with robust folding patterns are more likely to accommodate mutations without losing function, which in turn enhances their evolvability. By understanding the interplay between foldability and evolvability, as explored in studies on protein fitness landscapes, we can better predict which regions of a protein are most likely to contribute to resistance and tailor drug design accordingly^30^.

Despite these promising advances, several challenges remain. The complexity of bacterial evolution means that even highly constrained regions like our identified DHPS substructure, SB3 may eventually adapt through as-yet-undiscovered pathways. Additionally, the practical translation of these findings into effective drugs requires further research to identify small molecules that can specifically target such regions without causing off-target effects. Moreover, the dynamic nature of protein landscapes means that inhibitors must be designed with a deep understanding of both the current structure and potential evolutionary trajectories of the target protein.

Future research should focus on expanding the integrated evolutionary and structural approach to other clinically relevant proteins, exploring how these concepts can be applied across different pathogens. Additionally, continued exploration of protein fitness landscapes, particularly in identifying evolvability-enhancing mutations, will be critical for refining strategies to mitigate resistance. As antibiotic resistance continues to rise, innovative approaches that incorporate evolutionary constraints into drug design will be essential for sustaining the effectiveness of existing therapies and developing new ones. Ultimately, the identification of evolution-resistant regions across a range of proteins could provide a unifying strategy in the global effort to curb antibiotic resistance, making these findings a critical contribution to this ongoing challenge.

The integration of evolutionary dynamics with structural analysis offers a powerful framework for identifying new drug targets within key bacterial proteins. The identification of SB3 as an evolution-resistant, allosteric region in DHPS highlights the potential for developing next-generation antibiotics that remain effective even as pathogens attempt to evolve resistance. However, realizing the full potential of this approach will require addressing key challenges, including the development of specific inhibitors, understanding the long-term evolutionary dynamics of target proteins, and expanding this strategy to other critical enzymes involved in antibiotic resistance.

## Materials and Methods

### Generation of Sequence Space

We retrieved the sequence of *folP* gene encoding the dihydropteroate synthase of *Staphylococcus aureus* (SaDHPS) from UniProtKB (Uniprot ID - O05701) and used this sequence was used as the query for searching non-redundant databases in the BlastP. We obtained 1000 similar, non-redundant proteins sequences which were subsequently subjected to filtering at 75% sequence identity and query coverage. The dataset was then used to perform multiple sequence alignment (MSA)^31,32^ using Clustal Omega and which was further refined using AliView. Here we removed the sequences with a surplus of 20% gaps. This refinement prevented any bias from creeping into the MSA profile, which would have resulted in a contamination of the true and significant evolutionary signals. Post refinement, the MSA profile had 358 sequences with 278 positions. This MSA dataset signifies the sequence space and we used this for the sequence space analysis.

### Sequence Space Analysis

The sequence space was analyzed based on three broad aspects: conservation, co-variation and the mutational propensity of the residues comprising the sequence space. Conserved amino acid residues are characterized by little or no change across the evolutionary timescale. These residues are usually involved in key structural and functional roles. To characterize conserved positions, we used Shannon Entropy (SE), a concept in information theory, which measures the ‘uncertainty’ associated with/confined in a random variable. SE finds wide usage in evolutionary studies and is used in the form given below,

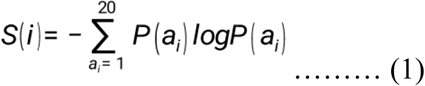

In the equation (1), ‘*i*’ represents the *i^th^* position in the sequence space, *S(i)* is the Shannon entropy value of the position ‘*i*’, *P(a_i_)* is the probability of amino acid ‘*a*’ to be present at the *i^th^* position and {*logP(a_i_)*} is the amount of information contained in a certain outcome of the random variable a_i_. With the decrease in SE values, the level of ‘uncertainty’ in finding a particular amino acid at a particular position decreases, which indicates the propensity of that amino acid residue to be more conserved. Higher SE values indicate a rise in the level of ‘uncertainty’ or more varying residues. The SE values were plotted against the position numbers for visualization.

While conserved residues are mostly linked with functional roles like the involvement in the catalysis of the enzyme, interaction sets between residues are mostly attributed to the preservation of the structural scaffold. We aim to study the conservation of such interaction networks by the investigation of the co-varying relationships of positions in the sequence space. Co-evolution/Co-variation proceeds in such a way that it couples the variation of positions to conserve the interactions which are crucial for the structural and functional integrity by contact formation. So, a study of co-varying positions reveals the co-evolution propensity of the positions, which makes the positions evolutionarily ‘Interdependent’ on each other. This interdependency brings a constraint in their evolution/variation, which is instrumental in preserving the overall architecture and in turn the function of the protein. We performed the Mutual Information analysis (MI analysis) to identify the positional correlations/constraints in the sequence space of DHPS. It is used in the form given below,

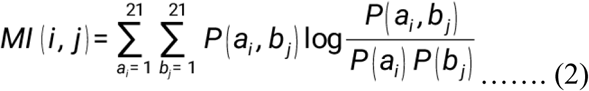

In the equation (2), *MI(i, j)* is the mutual information values, i.e. the co-evolution/co-variation propensity of *i^th^* and *j^th^* positions in the sequence space, *P(a_i_,b_j_)* indicates the probability of finding the amino acids ‘*a*’ and ‘*b*’ at the respective positions ‘*i*’ and ‘*j*’ simultaneously. The summation was done over the 20 amino acids and the gaps were treated as the 21^st^ amino acid. With the increase in MI values, the certainty of finding residue ‘a’ and *b*’ simultaneously at the *i^th^* and *j^th^*position increases, *i.e.,* the co-variation propensity increases and vice versa. To emphasize on the higher co-evolutionary signals in SaDHPS, we considered the top 50% of the co-varying pairs (cut-off value for MI = 0.43). Of these interdependent/co-varying positions/pairs, in order identify the co-varying pairs within the interaction distances, we considered only those positions, which were greater than 2 amino acids away on either side of/in the sequence space and were at less than 10 Å distance in the 3D structure of the protein. Using these pairs, we generated a co-evolutionary matrix and represented it as a heatmap. We generated the co-evolutionary network using the program Gephi where we applied Thomas Fruchterman & Edward Reingold graph layout^33^. It simulates the graph as a system of particles in a force directed layout. In the presented layout, the edges represent the position-position MI values and the nodes describe the residue positions. Here the edge width is defined by the MI values associated with interdependent pairs.

A common way of bringing about variation (or/and co-variation) in amino acids of proteins is via mutations. To unravel this aspect, we resorted to an unsupervised method for predicting the effects of point mutational effects both the positional residue information and inter-residue coupling effects. We performed a deep mutational scan to assess the quantitative effects of mutations and thus generated the mutational landscape for SaDHPS. By using the following equation,

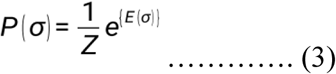

Where *σ* is a sequence and *P(σ)* measures the probability of the occurrence of that sequence in evolutionary timescale. *Z* is used to normalize the distribution. *E(σ)* is the evolutionary fitness energy analogous to the Gibbs free energy in statistical physics and used in the following form,

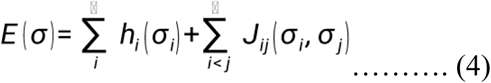

Here, *J_ij_* is a measure of the coupling strength between two positions *i* and *j* indicating the co-dependencies and *h_i_* is the site wise bias term. Contradictory to its analogy towards Gibbs free energy, the increase in *E(σ)* values indicate a higher evolutionary fitness and vice versa. The values of *E(σ)* were plotted as a heatmap for visualization.

### Structure Analysis

#### Model Selection

We retrieved the crystal structure of apo form of SaDHPS (1AD1) from RCSB and used the Chain A of the structure for further analysis^15^. In order to study the interactions of SaDHPS with substrates and the corresponding conformational changes in detail, we used the DHPt analogue bound form (1AD4)^15^ and the transition state analogue bound form (6CLV)^23^.

#### Structure Network Analysis

The structure network of a protein is the representation of the topological analysis of the complex 3D structure irrespective of its secondary structure and the folding type^34^. The correlated motions of the amino acid residues are significant for the conformational dynamics which bring about the functional activity of a protein. Normal Mode Analysis (NMA) is a computationally inexpensive method to study such dynamics, which finds the modes of oscillations close to the native state of the protein. Since the large-scale deformations associated with the higher modes of oscillations cannot really be tolerated by biomolecules, NMA provides an understanding for the functional dynamics of proteins. NMA indicates/deciphers the oscillations of the residues constituting the SaDHPS. These predicted motions were subjected to correlation analysis by the subsequent generation of a dynamic cross-correlation matrix (DCCM). Using the DCCM we generated the structure network of the apo form of SaDHPS (1AD1).

The generated structure network is a visual representation of the correlated functional motions of all residues of SaDHPS. Here the edges link the correlated/interacting residues (represented as nodes). The edge width indicates the magnitude of the correlation. To understand the structure/structure network in more detail, we split the all-residue network into a highly correlated coarse-grained community cluster network by using the Girvan-Newman clustering method. This led to the formation of discrete clusters of residues, or structure blocks (SBs) where the highly interacting residues are positioned together.

#### Frustration Calculation

We applied frustration analysis algorithm^26^ on the monomeric apo form of SaDHPS (PDB code: 1AD1, chain A) to localize and quantify energetic frustration in the protein structure in term of frustration index. The frustration index is a quantitative measure to assess the energetic compatibility of interactions within a protein structure. It measures how favorable or unfavorable specific interactions within a protein are, categorizing them as minimally, neutrally, or highly frustrated to reveal regions critical for structural stability, flexibility, and functional dynamics. The study identified areas of high frustration within the protein, indicating regions of energetic conflict important for its conformational dynamics and functional activity.

#### Combining Sequence and Structure Information

To integrate the sequence space as well as structural information of the significant regions of SaDHPS, we used two subsets of sequence-based and structure-based properties which encode the sequence space and structure space information, respectively. For this analysi, we considered the significant SBs (SB 1, 2, 3, 4, 5, 7 and 10) (explanation in results section). The sequence traits imposed were conservation index and coevolution index while the structural properties include mean hydrophobicity and order index.

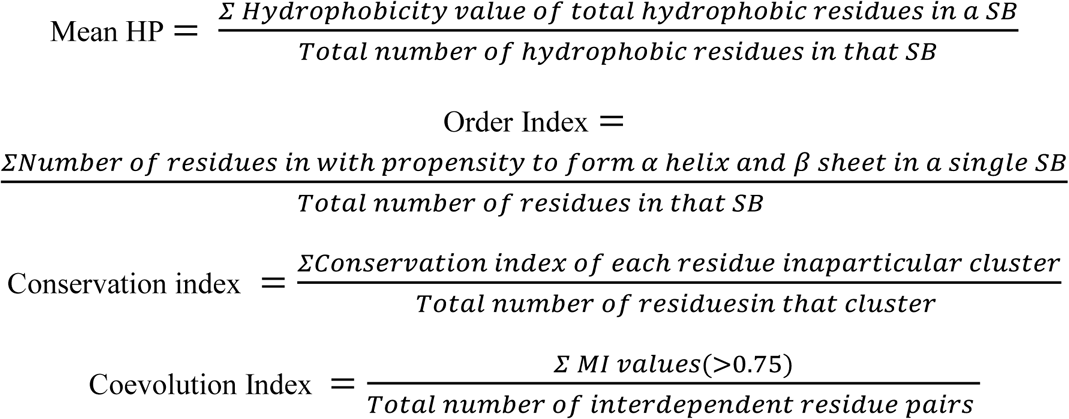

#### Pareto Optimality Analysis

We generated the dataset with the above mentioned indices as an ensemble of structural and evolutionary features: Mean HP, HP Abundance, Order Index, Conservation Index, and MI Index. Data pre-processing involved standardizing these indices via normalization, ensuring comparability across different scales and dimensions. This step was crucial for the accurate application of the subsequent Pareto optimality and utility function analyses.

Pareto optimality was employed to identify data points that exhibited optimal characteristics across all features^34^. Mathematically, a data point x in our dataset is Pareto optimal if there does not exist another point y such that y is at least as good as x in all indices and strictly better in at least one.

#### Utility Function Development

To further quantify the significance of each data point, a utility function U(x) was developed:

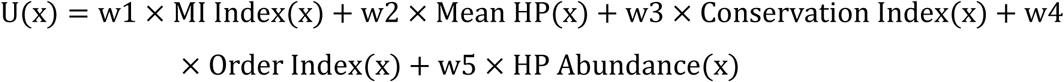

where *w_1_*,*w*2,*w*3,*w*4, and *w*5 are the weights assigned based on the normalized contributions of each feature to the Pareto front. These weights were calculated using the mean values of the features for the Pareto optimal points, normalized to sum up to 1.

#### Critical Features and SB Identification

To detect which features and Structure Blocks (SBs) were critical in determining Pareto optimality, we employed a two-pronged approach:

##### Feature Analysis

Each feature was evaluated for its impact on the Pareto front. We computed the mean of each feature for the Pareto optimal points and normalized these values to establish a relative importance scale. This scale was crucial in understanding which features had the most influence on defining Pareto optimality.

##### Utility Score Calculation for SBs

Each SB was assigned a utility score based on its feature values and the derived weights. The utility function, as previously defined, was applied to calculate these scores. Higher scores indicated a greater alignment with the Pareto optimal characteristics.

##### Ranking and Selection of SBs

SBs were then ranked based on their utility scores. The top-ranking SBs were identified as critical, given their high utility scores, which signified their significant role in the dataset’s Pareto optimality. This ranking provided a clear hierarchy of SBs in terms of their contribution to achieving optimal outcomes.

##### Cross-Feature Correlation Analysis

Additionally, a cross-feature correlation analysis was conducted to understand the interplay between different features within the top-ranking SBs. This analysis helped in identifying any synergistic or antagonistic relationships between features that contributed to the SBs’ high utility scores.

#### Principal Component Analysis

Principal component analysis (PCA) is a process of dimensionality reduction, where multidimensional data is reduced to a few principal components, while preserving the information (variance) as much as possible^34^. We deployed PCA to visualize the similarity or uniqueness of each of the shortlisted SB in the light of their sequence-based and structure-based properties defined earlier. Then disintegrated the sequence-based indices and structural indices separately into principal sequence component and principal structure component respectively, while retaining a high percentage of variance for each of the cases (∼100%). Then we projected the data with the reduced dimensions on to a hybrid sequence-structure 2-D subspace comprising of the principal components that were calculated.

### Druggability Analysis

#### Binding site validation

We investigated the potential druggability of the identified evolutionarily and structurally significant SB (SB3) by an unbiased, protein-wide search for small molecule or drug like molecule binding sites, based upon certain physical and biochemical properties. A binding site/pocket is generally a solvent accessible concavity on the surface of proteins. We used four different computational algorithms, each belonging to either computational geometry (Voronoi tessellation, Delaunay triangulation and alpha spheres) or protein threading methods to find out binding sites on the surface of SaDHPS. Also considered various biochemical and biophysical parameters while ranking the identified pockets, in terms of their capabilities of binding a small molecule or specifically a drug molecule. These parameters were either solvent-accessible volume, a composite Druggability score, a mathematically defined confidence score or a z score computed from the RMSDs and Blosum62 matrices between superimposed residues. Thus, by combining these various methods for pocket finding and ranking, minimized the limitation or bias of a particular method. We selected the best ranking pockets with respect to small molecule binding and listed the residues falling under the SB (SB3) to find out the small molecule binding capabilities of the region. To assess the mutational sensitivities of the druggable residues of the SB, for every position, we calculated the mean of the 19 fitness scores upon substitution, obtained from the deep mutational scan. We plotted the mean fitness scores of the 13 druggable residues in SB3 and marked the median of the distribution in the plot.

## Acknowledgement

DS and KC acknowledge the Director, IICB. AS, AC and SC acknowledge the Director, Birla Institute of Technology and Science-Pilani, India.

## Author Contributions

SC and KC planned the overall project outline. DS and SC designed the work plans and performed the computational experiments and analyses. VU critically analyzed the manuscript and performed the comparison of distributions of MDP, RMSF, and Shannon entropy. DS and AS wrote the manuscript, which was then refined by SC and KC. All authors approved the final manuscript.

## Declaration

The authors do not have any conflicts of interest.

## Supplemental information

### Supplemental Figures

**Supplemental Figure S1:**
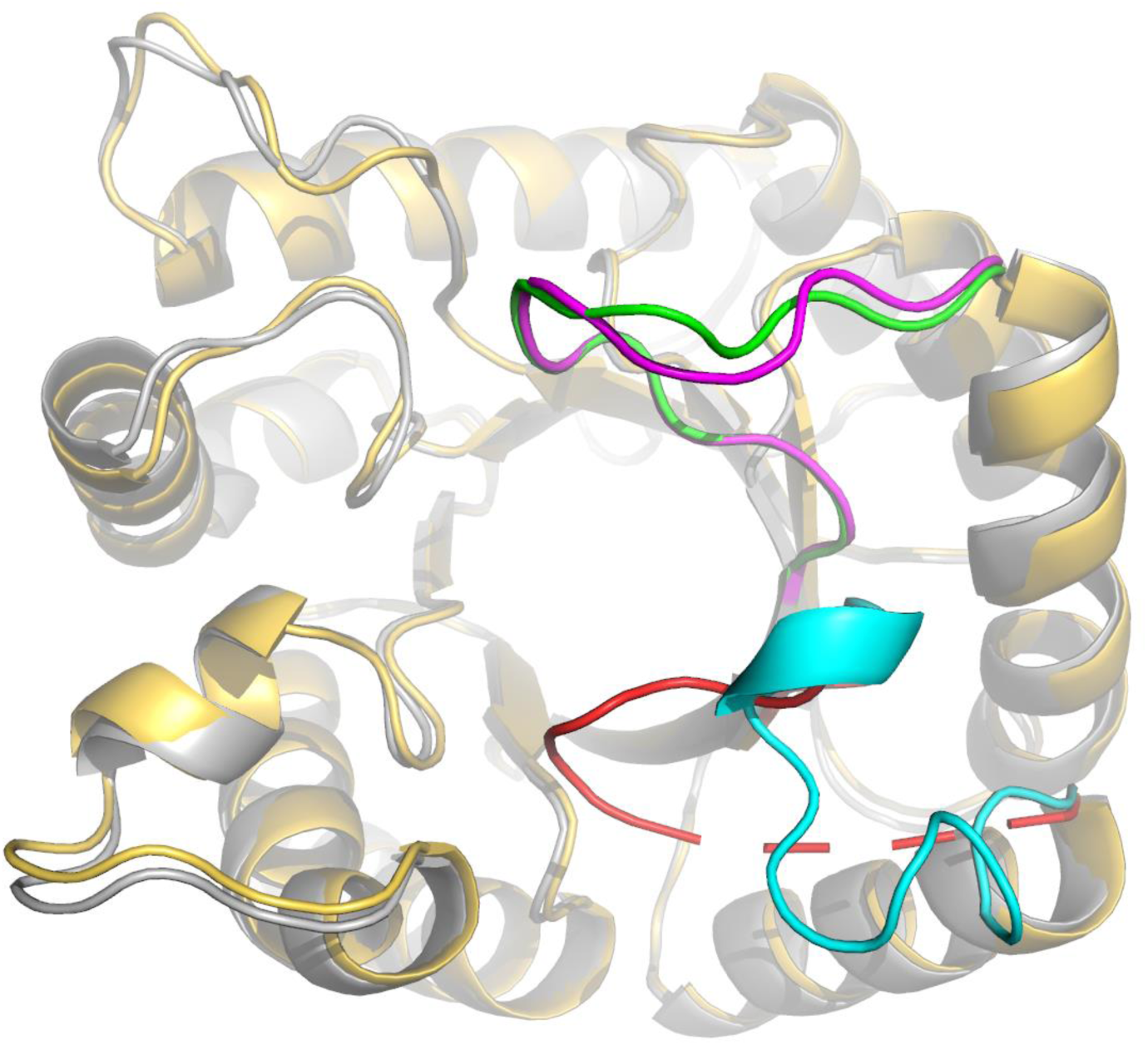
Superposition of apo (1AD1, gray) and transition state analogue (6CLV, golden) bound structure of SaDHPS. Loop I is colored cyan in the apo structure and red in the transition state structure. Similarly, loop II is colored magenta in the apo structure and green in the transition state structure. These two loops move (specially Loop I has larger movement) towards the active site in the transition state of the bound structure (red color) as compared with the apo form of the enzyme (cyan color).

**Supplemental Figure S2:**
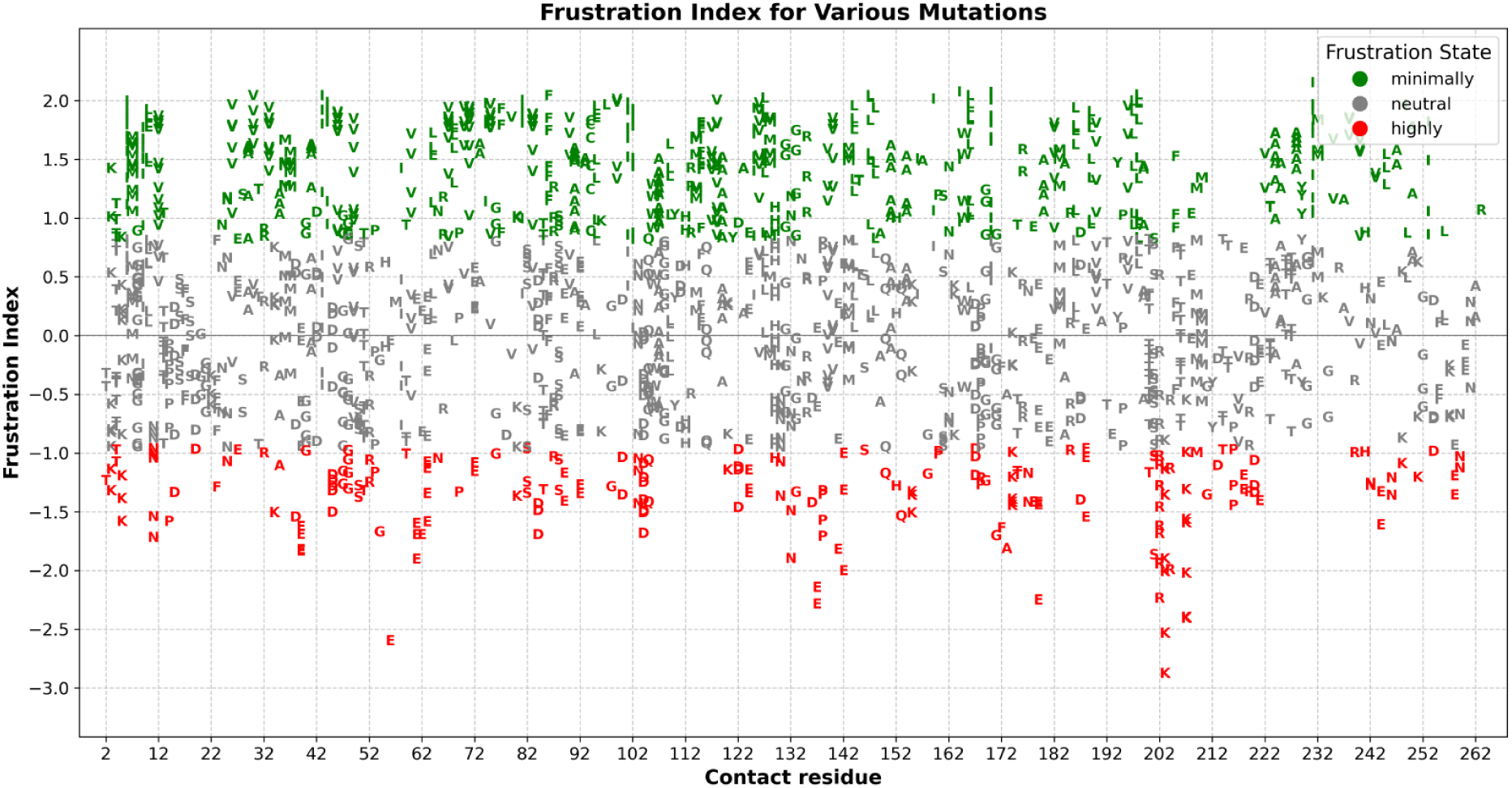
The frustration index values for residue interactions, which are based on variations in protein mutational energy.

### Supplemental Tables

**Supplemental Table 1:**
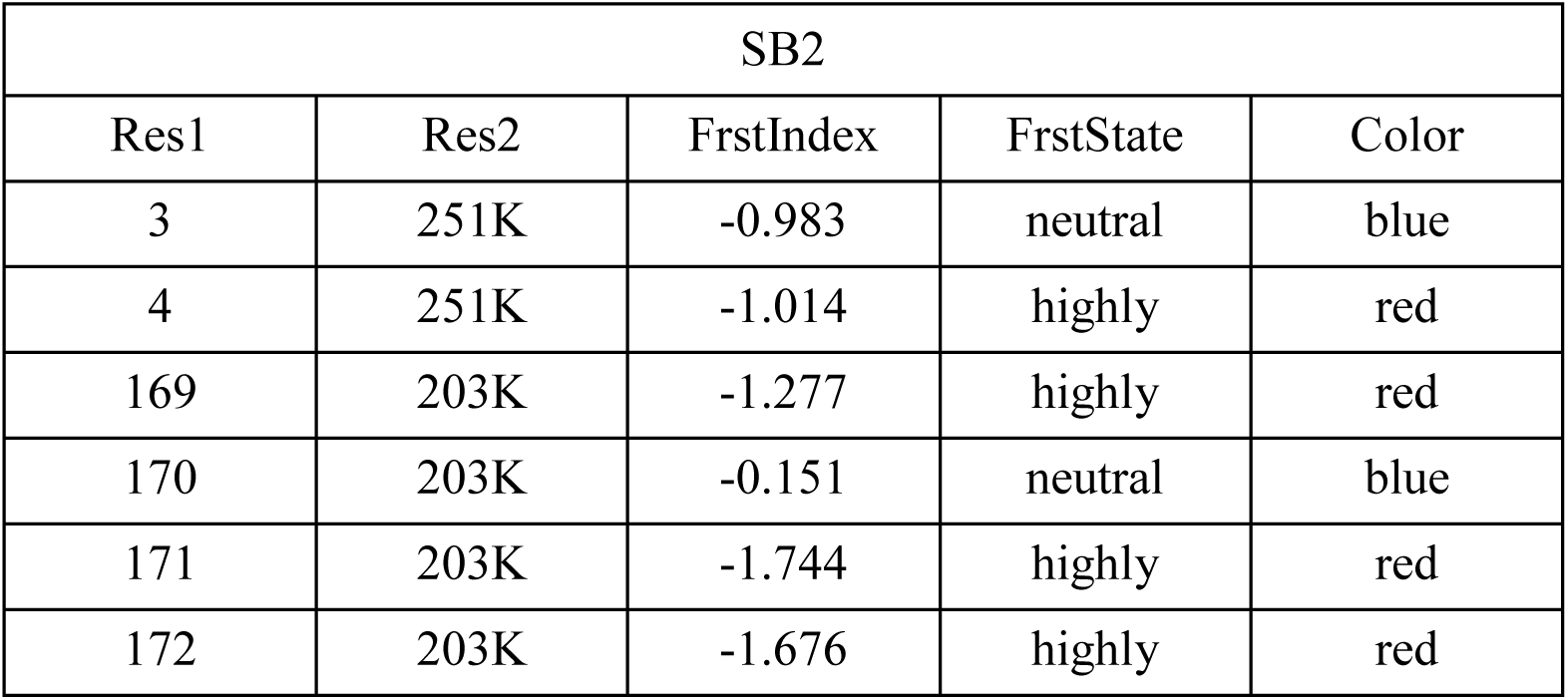
Frustration index results of SB 2. The table shows the interaction of SB 2 with the Dimer interface region (251) and (203), which is interaction with both DHPt and pABA.

| SB2 |  |  |  |  |
| --- | --- | --- | --- | --- |
| Res1 | Res2 | FrstIndex | FrstState | Color |
| 3 | 251K | -0.983 | neutral | blue |
| 4 | 251K | -1.014 | highly | red |
| 169 | 203K | -1.277 | highly | red |
| 170 | 203K | -0.151 | neutral | blue |
| 171 | 203K | -1.744 | highly | red |
| 172 | 203K | -1.676 | highly | red |

**Supplemental Table 2:** Frustration index results of SB 3. The table shows the interaction of SB 3 with Loop I (11, 12), Loop II (47, 48), and dimer interface region (244, 251).

| SB3 |  |  |  |  |
| --- | --- | --- | --- | --- |
| Res1 | Res2 | FrstIndex | FrstState | Color |
| 6 | 244E | 0.518 | neutral | blue |
| 6 | 251K | 0.635 | neutral | blue |
| 9 | 11N | 0.199 | neutral | blue |
| 9 | 47G | 0.413 | neutral | blue |
| 10 | 12V | 1.812 | minimally | green |
| 10 | 47G | 0.77 | neutral | blue |
| 10 | 48G | 0.692 | neutral | blue |
| 35 | 244E | -0.721 | neutral | blue |
| 36 | 244E | 0.245 | neutral | blue |
| 38 | 244E | -1.583 | highly | red |
| 39 | 244E | -1.862 | highly | red |
| 40 | 244E | -0.95 | neutral | blue |
| 40 | 251K | -0.896 | neutral | blue |
| 41 | 244E | 0.352 | neutral | blue |
| 45 | 47G | -1.225 | highly | red |
| 46 | 48G | 0.845 | minimally | green |
| 242 | 244E | -1.313 | highly | red |

## Notes

### Competing Interest Statement

The authors have declared no competing interest.

